# SARS-CoV-2 Omicron Variant Wave in India: Advent, Phylogeny and Evolution

**DOI:** 10.1101/2022.05.14.491911

**Authors:** Urvashi B. Singh, Sushanta Deb, Rama Chaudhry, Kiran Bala, Lata Rani, Ritu Gupta, Lata Kumari, Jawed Ahmed, Sudesh Gaurav, Sowjanya Perumalla, Md. Nizam, Anwita Mishra, J. Stephenraj, Jyoti Shukla, Deepika Bhardwaj, Jamshed Nayer, Praveen Aggarwal, Madhulika Kabra, Vineet Ahuja, Subrata Sinha, Randeep Guleria

## Abstract

SARS-CoV-2 evolution has continued to generate variants, responsible for new pandemic waves locally and globally. Varying disease presentation and severity has been ascribed to inherent variant characteristics and vaccine immunity. This study analyzed genomic data from 305 whole genome sequences from SARS-CoV-2 patients before and through the third wave in India. Delta variant was responsible for disease in patients without comorbidity(97%), while Omicron BA.2 caused disease primarily in those with comorbidity(77%). Tissue adaptation studies brought forth higher propensity of Omicron variants to bronchial tissue than lung, contrary to observation in Delta variants from Delhi. Study of codon usage pattern distinguished the prevalent variants, clustering them separately, Omicron BA.2 isolated in February grouped away from December strains, and all BA.2 after December acquired a new mutation S959P in ORF1b (44.3% of BA.2 in the study) indicating ongoing evolution. Loss of critical spike mutations in Omicron BA.2 and gain of immune evasion mutations including G142D, reported in Delta but absent in BA.1, and S371F instead of S371L in BA.1 could possibly be due to evolutionary trade-off and explain very brief period of BA.1 in December 2021, followed by complete replacement by BA.2.

## Introduction

The origin of SARS-CoV-2 has been ascribed to a recombination in Bats, enriching it with the capability to utilize enzyme furin from human cells, enabling the spike protein to bind the Neuropilin1 on human cells, hence facilitating cellular entry and replication. The virus was thus different from SARS and MERS. The S:D614G mutation improved its ability to bind to ACE2 receptors, further enabling the ease of infecting humans, and thus widespread transmission ^1^. The H69/V70 deletion enabled increase in infectivity by two-fold and the N501Y and K417N enabled stronger binding with ACE2 receptors ^2^, the latter along with E484K mutation were shown to enable reduction in neutralization by antibodies ^3^.

WHO monitors the evolution of SARS-CoV-2 and evaluates if specific mutations affect the virus behavior. The Omicron variant (B.1.1.529) was reported on 24 November 2021, to WHO from South Africa. This variant was reported to have a large number of mutations, and increased risk of reinfection. The variant spread to all continents and was responsible for a large number of cases globally. It was detected at faster rates than previous surges and was designated a variant of concern. Countries were advised to enhance surveillance and sequencing efforts, field investigations to understand impact of this variant on COVID-19 epidemiology, disease severity, immune response, virulence, transmissibility, immune, diagnostic or therapeutic escape ^4^. In February 2022, the new sub-lineage BA.2 was reported to be spreading widely. WHO’s Technical Advisory Group on SARS-CoV-2 Virus Evolution recommended that this sub-lineage should continue to be considered a variant of concern, and remain classified as Omicron, based on data available on its severity, reinfection capability, diagnostics ^4^.

The Omicron variant continues to be the dominant variant globally, including sub-lineages BA.1, BA.1.1 (or Nextstrain clade 21K) and BA.2 (or Nextstrain clade 21L). BA.2 differs from BA.1 in spike and other proteins amino acids and has been shown to have growth advantage over BA.1 ^4^. BA.2 appears to be more transmissible than BA.1, and hence is the most reported Omicron sub-lineage. Animal studies from Japan have highlighted that BA.2 may cause more severe disease in hamsters, though this was not reported from vaccinated population in UK, Denmark and South Africa. Studies have also demonstrated strong protection against reinfection with BA.2 following infection with BA.1 ^5^. In India, Omicron transmission continued through December 2021-February 2022. December 2021 strains were BA.1 to begin with, but soon BA.2 replaced as the predominant strain. Most cases were reported as mild, attributable to reduced severity, infrequent lung involvement and higher population immunity. Higher number of cases indicating higher transmission resulted in more hospital admissions. Deaths were primarily reported in unvaccinated population ^6^.

Our study was designed to understand the genomic differences and evolutionary changes in the variants prevalent in and around the city of Delhi, before and during the third wave of SARS CoV-2.

## Methods

### Sample Processing

Samples were collected as part of routine testing for patients reporting to the Emergency Department at the All India Institute of Medical Sciences, with symptoms of high grade fever, shortness of breath and headache. The samples were subjected to whole-genome sequencing following the IEC approval (IEC-679/03.07.2020,RP-32/2020).

Nasopharyngeal and oro-pharyngeal swabs were collected from individuals presenting to Emergency Department, and tested using Cartridge based Nucleic Acid Amplification Test (CBNAAT) in the Biosafety Level-3 laboratory, Department of Microbiology from June 2021 to March 2022. Patients presenting with respiratory symptoms suggestive of COVID-19, and patients planned for emergency interventions were enrolled in the study. Sample was collected as per standard procedures using a nylon-flocked swab and transported to the laboratory in Viral Transport medium (TrueNat uses Viral lysis buffer). Samples were subjected to CBNAAT in the BSL-3 laboratory (either GeneXpert or TrueNat) ^7,8^. Xpert Xpress SARS-CoV-2 assay detects envelope (*E*), and nucleocapsid (*N2*) genes of SARS-CoV2 with sample processing control.

### Next generation sequencing

Nasopharyngeal and throat swab samples from COVID-19 cases, with Ct value ≤ 25 were processed for viral genome sequencing using COVIDSeq Assay (Illumina Inc, USA) on MiSeq sequencing platform using the manufacturer’s protocol. Viral RNA was isolated from the 140 μl of viral transport media containing l of viral transport media containing swabs using QIAamp Viral RNA Mini kit (Qiagen, Hilden, Germany) following the manufacturer’s instructions. Briefly, cDNA synthesized from RNA samples was subsequently amplified using a primer pool capable of amplifying 98 targets across SARS-CoV-2 genome. The amplified products were tagmented and ligated with adaptors using Nextera UD

Indexes Set A, followed by enrichment of desired fragments. The samples were pooled and then quantified using DNA HS assay kit Qubit 4.0 fluorometer (Invitrogen Inc. USA). The fragment sizes of the pooled libraries were assessed with DNA HS Kit (Agilent Technologies, Santa Clara, USA) on Agilent Bioanalyser. The pooled library was further normalized to 4nM concentration, denatured and loaded onto a Miseq V3 Flow Cell (150 cycles) to carry out paired end sequencing with a read length of 2×75 on MiSeq platform. The FASTQ files were aligned against the SARS-CoV-2 reference genome (NC_045512.2) ^9^. The FASTA files were generated and analysed using Illumina DRAGEN COVID Pipeline and DRAGEN COVID Lineage Tools (v3.5.3).

### SARS-CoV-2 lineage assignment

The complete genome sequences of 305 SARS-CoV-2 strains isolated from June-2021 to February -2022 in Delhi, India were used throughout this study for phylogeny and codon usage analysis. The phylogenetic analysis was performed to get an insight on global and local genetic diversity of the SARS-CoV-2 variants considered in this study. The lineages of viral strains were identified using Pangolin web service ^10^. The accession numbers and other metadata about the isolates can be found at Supplementary Table S1.

### Phylogeny and mutational analysis

The SARS-CoV-2 genome sequences obtained from this study were subjected to maximum likelihood (ML) phylogenetic analysis using IQ-TREE v2.1.2 ^11^. The whole-genome sequences were aligned using were aligned using MAFFT v7.475 ^12^, the aligned sequences were further used as input for IQ-TREE program build in nextstrain pipeline and downstream analysis steps were followed to generate the phylogenetic tree. Generated tree file were visualized using the FigTree program v1.4.4 (available at: http://tree.bio.ed.ac.uk/software/figtree/). To determine the phylogenetic clade, isolated SARS-CoV-2 genomes were compared against the global Nextstrain database (https://nextstrain.org/ncov/global) using the combined package of Augur, the MAFFT and the IQ-tree software embedded in Nextstrain pipeline. The nucleotide and amino acid substitution analysis were performed by augur pipeline and other accessory module of nextstrain pipeline ^13^.

### Codon usage analysis

The codon usage analysis was performed using Bash and R scripts of CUBES package ^14^. The modal codon usages ^15^ were calculated for the genes from given viral genomes with the help of calculate_modals2.pl script implemented in CUBES package (https://github.com/maurijlozano/CUBES).The correspondance analysis (CA) ^16^ based on relative synonymous codon usages (RSCUs) of viral genomes were performed by CodonW Program ^17^. Raw-codon-count (RCC) and RSCU based correspondance analysis (CA) were calculated using coa_Ccounts_GNM.sh and coa.sh scripts of CUBES package ^14^.

### Analysis of tissue-specific adaptation

For the analysis of codon usage similarity extent of delta and Omicron variant with the tissue specific highly expressed genes, a recently introduced codon usage metric similarity index (SiD) or D(A,B) has been adopted in this study.

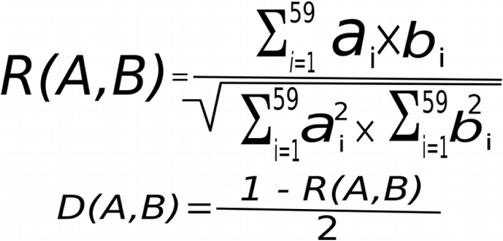

where R(A, B) represents cosine value between A and B special vectors, indicating the degree of similarity between two samples on the basis of the overall codon usage pattern. The a_i_ and b_i_ is defined as RSCU values of the specific codons in the samples selected for pairwise comparison. The D(A,B) represents influence of overall codon usage of tissue specific genes on Sars COV-2 variants. When D(A,B) value is closer to zero, higher similarity extent of synonymous codon usage patterns between two samples. The list of tissue specific highly express proteins were retrieved from the Human protein atlas database (https://www.proteinatlas.org/; Uhlén et al.,2015) ^18^. The coding sequences of those highly expressed proteins were extracted from transcript file (available in CAIcal server: http://genomes.urv.es/CAIcal/human_genes_from_ensembl; Puigbò et al) ^19^ of human genome using custom perl script.

## Results

### Study Population

Patients reporting to Emergency Department with symptoms suggestive of SARS-CoV-2 or those who were tested for SARS-CoV-2 prior to intervention/procedure and detected positive for SARS-CoV-2 from June 2021 to February 2022 were enrolled. This duration covered the period before and during the third wave in Delhi during the month of January 2022. Of 305 samples sequenced, 16.7% were in age group 0-18, 40.6 % in 19-40 years and 42.6% in 41-80 years, with a mean age of 37 and 41.6% females. Among them 31.6% had received vaccination and 68.5% patients had some comorbidity (9.6% chronic pulmonary disease, 26.3% cancer, 14.4% hypertension, 7.7% chronic kidney, 10% cardiovascular ailment, 11.5% Diabetes mellitus, 9% neurological disease, 3.8% hypothyroidism and 22% less severe health issues). Of those with comorbidities, 62% were admitted while the remaining advised home isolation. Death was reported in 3.3% study population. Delta variants affected patients without any comorbidity in 97% (30/31), BA.1 affected 50% patients without comorbidity, BA.2 lead to disease in 23% patients without comorbidity, while 77% patients had Diabetes, hypertension, or some other comorbidity. (Table S1).

### Phylogeny and lineage distribution

The SARS-COV-2 variant circulating in Delhi was predominantly Delta variant during June 2021 and was replaced by Omicron BA.1 in December 2021, which stayed a very short duration, started with very few cases in mid Dec-2021 and continued till the first half of January 2022. Unlike BA.1, Omicron BA.2 lineage was predominant in Jan-Feb, 2022, interestingly we observed that the BA.2 variant replaced BA.1 in February 2022.The phylogenetic tree constructed from the whole genome sequences of study isolates clustered into three major clades. The clades colored in red, blue and pink are representing the Delta, Omicron BA.1 and Omicron BA.2 variants. In the present study among the total sequences, 26 % were Delta B.1.617.2 and AY.12, 4% Omicron BA.1 and 69% Omicron BA.2 lineage (Fig. S1). A closer look into the BA.2 clade reveals that Omicron BA.2 isolates collected in Feb 2022 are clustered in multiple distinct clades away from BA.2 isolates of December and January (Fig. S2). A time-scale phylogenetic tree constructed using Next-strain pipeline to identify the phyletic position of Delhi isolates in global context, our isolates are represented in clades with specific colored dots on global phylogenetic tree (Fig. 1).

**Figure 1:**
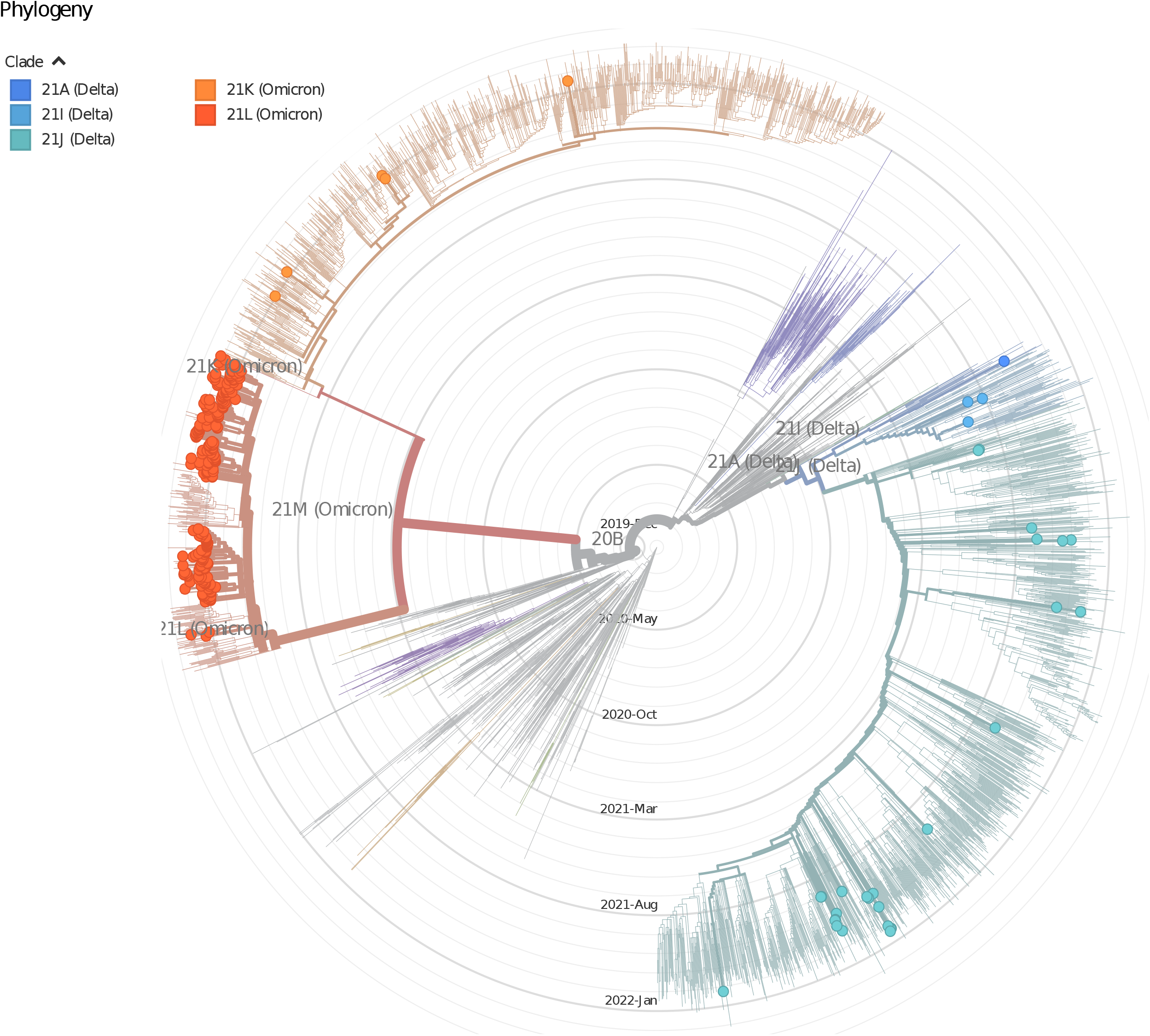
The time stepped phylogenetic tree (using Nextstrain pipeline)^13^ showing phyletic placement of SARS-CoV-2 variants of Delhi. The picture shows distribution of lineages of Delhi isolates, the nodes are colored according to clades.

### Tissue specific adaptation of isolated SARS-CoV-2 variants

In this study human tissues from two main anatomic sites (Lung and Upper respiratory tract) that are reported to harbor a high viral load during COVID-19 infection were selected for tissue specific adaptation analysis of SARS-CoV-2 variants. The similarity index (SiD) estimates similarity between host and pathogen codon usage profile, and has been exploited to measure viral adaptation to the host ^20^. The SiD value ranges from 0 to 1, if the value is close to 1 it indicates low viral adaptive capability inside the host system ^21^. In the present study it has been observed that SiD value of Delta variant is relatively lower for lung tissue, however unlike delta, Omicron shows lower SiD value for upper respiratory tract (Fig. 2).

**Figure 2:**
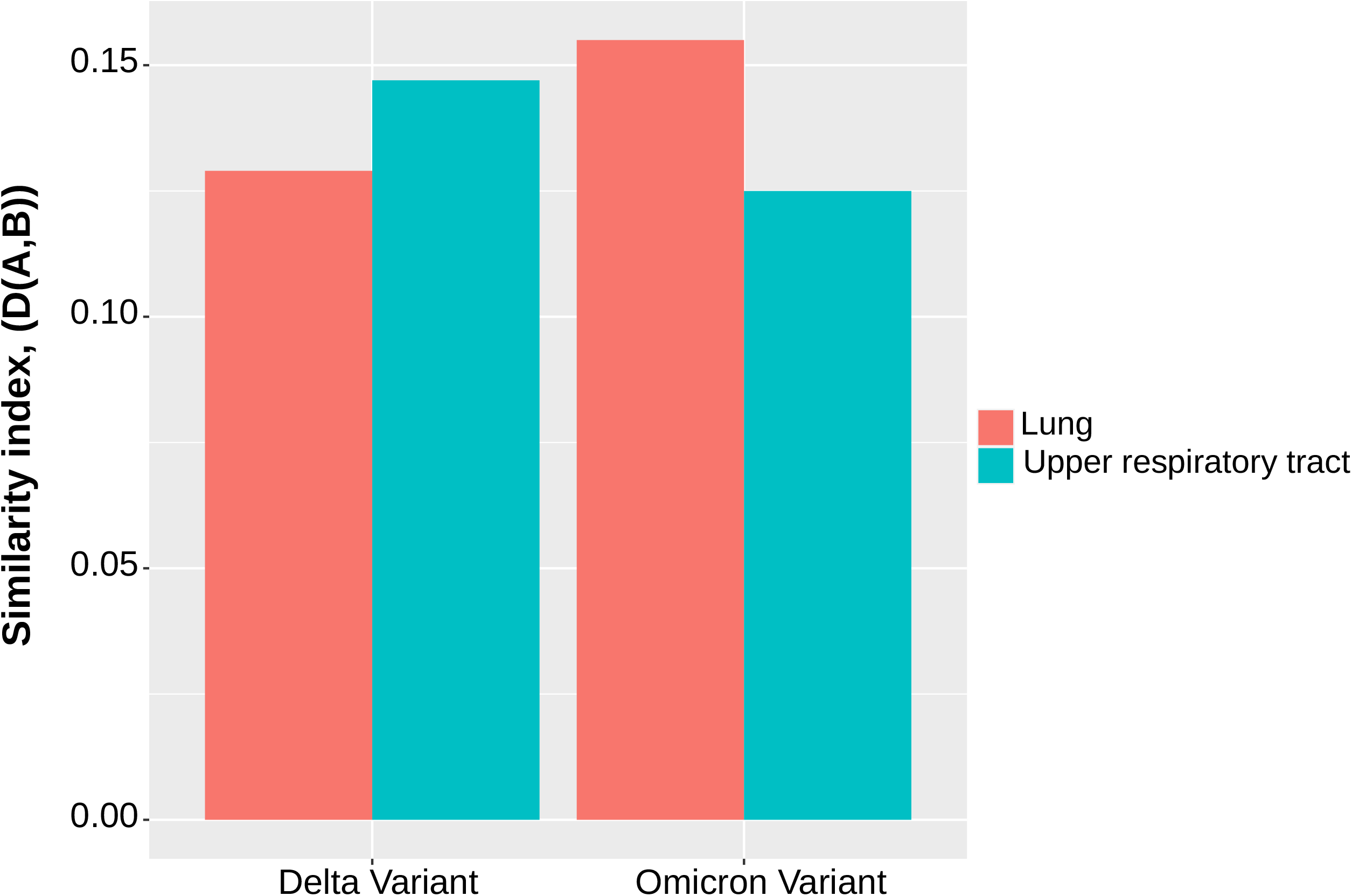
Similarity of codon adaptation index of Omicron and Delta genome. D(A, B)^20^ indicates the difference in codon usage similarity extent between two variants with the tissue specific high expression genes from two anatomic sites with high viral load.

### Varied mutation profile in Omicron and Delta variants of Delhi

A filtration criterion of (NNN= ≤ 1%) was applied to genomes for mutation analysis. After filtration, 31 Delta and 185 Omicron genomes were analyzed and found to have 1177 and 11488 amino acid substitutions in Delta and Omicron genomes respectively. Four substitutions are shared among all variants, two (T478K and D614G) in S glycoprotein and two (T3255I and P314L) in ORF1a and ORF1b, respectively (Table S2 and S3). Nearly 50% of substitutions belong to the Spike region in the Omicron genomes, 57% of total spike substitutions were confined to RBD (receptor binding domain) region. An average of 17.2 and 60.6 synonymous mutations were recorded in Delta and Omicron variants, respectively. Gene-specific amino acid mutation profiles of Delta and Omicron variants were compared (Fig. 3). Comparison of critical spike mutations was performed between BA.1 and BA.2 variants, three mutations (S371L, G446S and G496S) reported to be immune evasion mutations ^22,23^, were found in Omicron BA.1 but were absent from BA.2 variant, while three other immune evasion mutations were gained in 98% of BA.2 study isolates, T376A, D405N and R408S in the Spike region ^24^. Two mutations were gained (G142D and S371F) by BA.2 variant (Table S3). G142D was reported in Delta but lost in BA.1, while S371F is a better immune evasion substitution instead of S371L found in BA.1 variant ^25^. A new mutation S959P in ORF1b was found consistently in 44.3% BA.2 strains after December 2021. P314L (exists in linkage disequilibrium with D614G) ^26,27^, in ORF1b region, seems to be conserved in currently prevailing Delta and Omicron variants of Delhi. The median number of total mutations was 65/sample in Omicron BA.1 and 70/sample in BA.2 variants.(Fig. S3). Pattern of amino acid and nucleotide mutations in Delta and Omicron variants is depicted in Figure-3 and Figure-S4.

**Figure 3:**
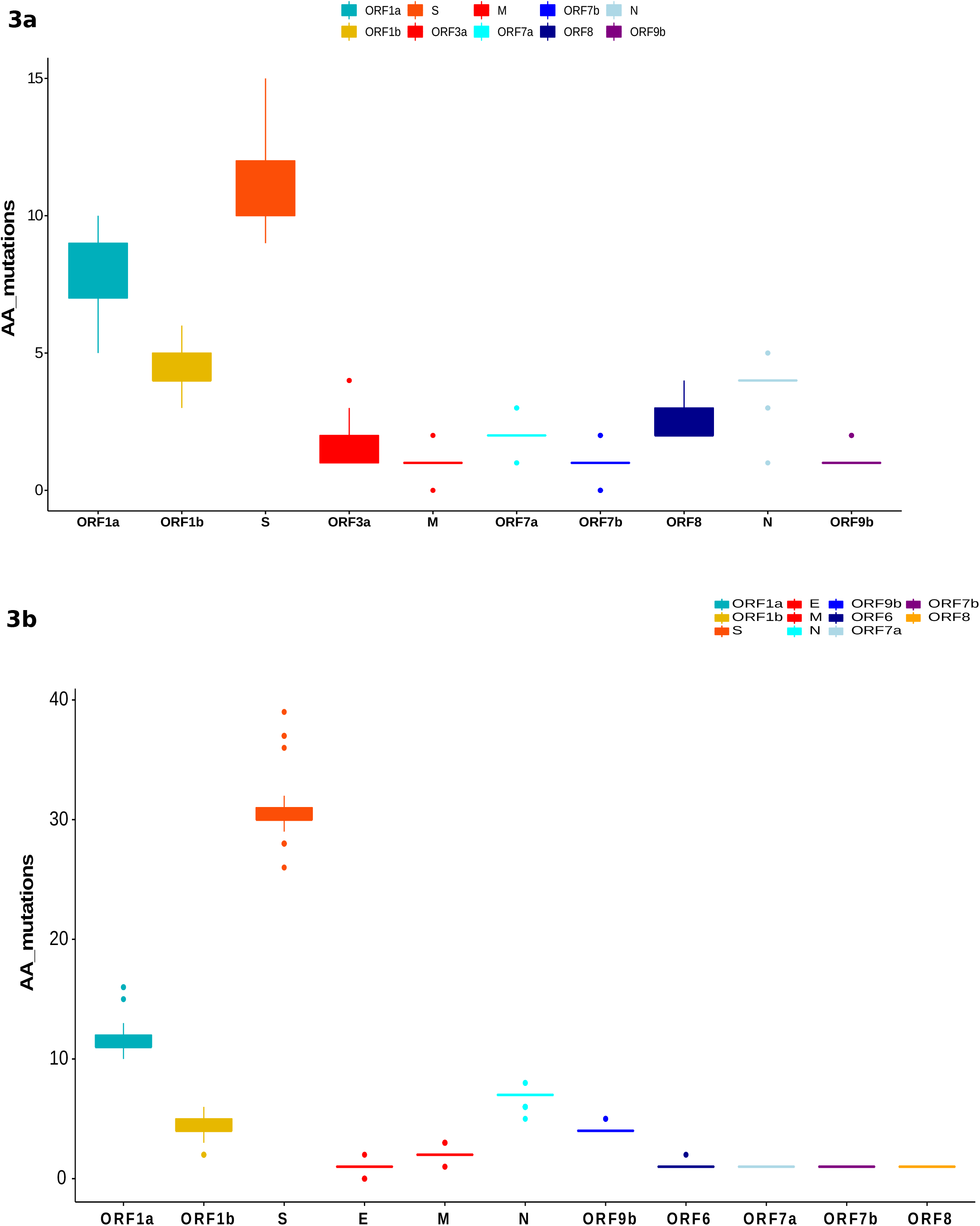
Amino acid substitutions in open reading frame (ORF) in Delta(3a) and Omicron(3b). Boxes contain the 25th and 75th percentiles, and whiskers show the minimum and maximum values. Thirty eight gene specific common substitutions in ORF1a (4), ORF1b (2), S (19), E (1), M (2), N (6) and ORF9b (4) were observed in Delhi isolates of Omicron BA.1 and BA.2 variants. BA.1 genomes harbored 24 signature mutations, in ORF1a (6), S (17) and M (1). BA.2 genomes contain 22 signature mutations, in ORF1a (7), ORF1b (2), S (11), ORF3a (1) and ORF6 (1).

### Variation in codon usage and selection of high-frequency codons

A comprehensive analysis of codon usage frequency was performed to understand the trends of evolution currently operating on the total genetic contents of the SARS-CoV-2 strains circulating in Delhi. A significant difference in codon usage among the genomes of SARS-CoV-2 variants was observed between Delhi isolates from June-2021 to February-2022 (Fig. 4,a1). The figure shows the Delta genome clustered (red) separately from the Omicron genome (yellow) (Fig. 4,a1), which indicates Delta variants are genetically distant from Omicron variants and a notable genetic distance was observed among the currently circulating Omicron variants of Delhi (Fig. 4,a2). However, unlike Omicron variant, not much genetic variability was observed among the Delta variants of Delhi. Few genomes were represented as a single dot (singleton) in figure 4, mainly due to the high ambiguity (Multiple Ns) in its nucleotide sequence. In addition it is evident from the analysis that the genomes of SARS CoV-2 variants were enriched in C and U ending codons (Fig. 4,b1-b3, Right graph).

**Figure 4:**
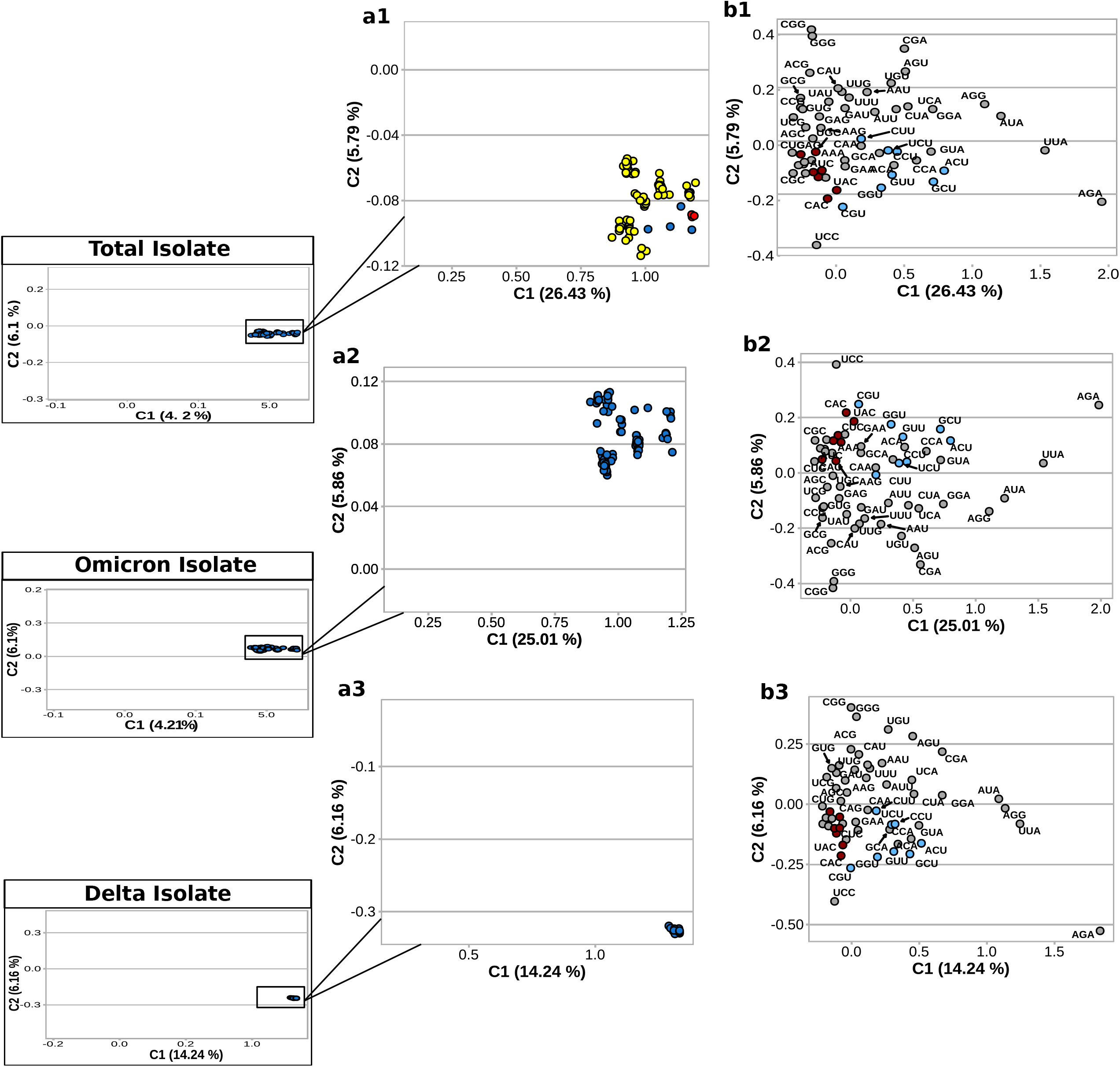
Correspondence Analysis (CA) using raw codon counts (RCC) plots the codon usage profiles of the genomes of SARS-CoV-2 variants from Delhi. Magnified view of the boxed region in left panel (A, B and C) of the figure displays position of the model codon usage in all genomes; in middle panel yellow and red clusters represent Omicron and Delta genomes (a1 graph). The graphs separating Omicron variant (a2 graph) and Delta variant (a3 graph) can be found at lower middle panel. The loading plots b1 (all isolates), b2 (Omicron isolates) and b3 (Delta isolates) describing codon relative weight in the first two principal-components of the CA presented in extreme right panel of the figure. Codons with the highest codon usage frequency (CUF) enrichment for each amino acid are colored dark brown and light blue for codons corresponding to C or U, respectively.

## Discussion

This study demonstrates tissue adaptation, whole genome codon usage pattern and evolution of the recently emerged Omicron variant and the Delta variant circulating in Delhi. Paz et al have reported codon usage based evolutionary insights into SARS-CoV-2 Omicron variant ^28^. Phylogenetic relationship among the SARS-CoV-2 variants of Delhi disclosed a distinct grouping of February isolates of Omicron BA.2 variants (Fig. 4), highlighting genetic diversity of February isolates from December and January. The genetic diversity in BA.2 isolates reflects on the codon usage plot. Multiple separate clusters of the Omicron genomes represent the genetic distance among the Omicron variants currently dominating in Delhi (Fig. 4, a2; left graph). The findings indicate that the Omicron variant in Delhi is under the process of genetic evolution, and could give rise to multiple Omicron variants in the near future.

Simmonds et al discussed C→U SNP, as being predominant in all SARS-CoV-2 variants. We found similar substitutions in Delta and Omicron in Delhi uniformly (Fig. 4, b1-b3; right graphs). The U ending codons were found approximately four times higher than other substitutions ^29,30^.

Several reports have ascribed lesser severity of Omicron variants to lower lung involvement in comparison to Delta variant and higher replication in bronchi ^31^. Our findings demonstrate that Omicron variant is more adaptable to the upper respiratory tract compared to the lung while Delta variant has relatively higher adaptability towards lung tissue. This further ascribes higher transmission capability to Omicron variant.

Studies report maximum amino acid substitutions in the spike region, specifically the RBD region in Omicron variants ^32,33^. Current study found similar results. Desingu et al reported that recent Omicron variants are losing the mutations found in the RBD region of BA.1. Interestingly, we found that BA.2 Omicron variant in Delhi has lost critical mutations in the spike region when compared to the BA.1 variant, including some previously reported immune evasion mutations, while gaining other immune evasion mutations ^34–36^. Delta contained lesser number of total mutations compared to Omicron variants in Delhi. Despite the loss of critical mutations in Spike protein, BA.2 acquired several mutations in ORF bringing the total mutations to 70/sample compared to the Wuhan strain, maximum reported so far ^32,35^.

Viral glycoproteins are known to be subjected to evolutionary trade-offs, mutations responsible for higher infectivity and transmissibility could have detrimental effect on another property, such as host immune escape capability ^37^. Weismann et al demonstrated enhanced susceptibility to neutralization due to D614G mutation in spike protein, the mutation that was shown to be responsible for higher transmission of SARS-CoV-2, indicating the evolutionary trade-offs operating on spike mutations in SARS-CoV-2 ^38^. The effect of a mutation on viral fitness could shape the fate of such mutations. Possibly, unfavorable mutations in early Omicron BA.1 variant could have led to loss of fitness, long term spread and survival, hence explaining loss/gain of critical mutations in Omicron BA.2 lineage, its wider and continuing spread and evolution.

## Supporting information

Four supplement figure and two supplement spread sheet

## AUTHOR CONTRIBUTIONS

UBS conceived and formulated the study, planned the work, and wrote the manuscript with SD, LR, LK, and DB. UBS, RC, KB, SS, RiG, MK, VA and RG contributed to diagnostics and laboratory man-agement. JA, SG, SP, MN, AM, JSt conducted diagnostic tests, LR conducted Whole Genome Se-quencing, SD conducted Bioinformatics. JN and PA provided clinical care. SD, DB, LK and JS con-tributed to the data analysis.

ACKNOWLEDGEMENTS

Authors acknowledge funding from All India Institute of Medical Sciences, New Delhi SS acknowledges the J C Bose Fellowship of the Department of Science and Technology, India.

## DATA AVAILABILITY

The sequence data is available in GISAID

SUPPLEMENTARY DATASETS

Supplementary Data 1: GISAID submission with Metadata

Supplementary Data 2: Mutations in genomes of Delta Variants from the study

Supplementary Data 3: Mutations in genomes of Omicron Variants from the study Supplementary Figures

Figure-S1 (Supplement):

Lineage distribution of SARS-CoV-2 genomes isolated from Delhi during the time period of June-2021 to Feb -2022.

Figure-S2 (Supplement):

Maximum likelihood phylogenetic tree (ML, GTR, 1,000 bootstrap) of the 305 genome sequences obtained in this study. The scale of the phylogenetic branches is based on nucleotide substitution per site.

Figure-S3 (Supplement):

The phylogenetic relationship and mutation burden across SARS-CoV-2 variants isolated from Delhi. The radial axis shows the total number of mutations in different lineages.

Figure -S4 (Supplement):

Nucleotide mutations in Omicron and Delta genomes. Boxes contain the 25th and 75th percentiles, and whiskers show the minimum and maximum values.

**Supplement Table S1**

Metadata of Delhi SARS-CoV2 isolates with GISAID Accession Number.

**Supplement Table S2**

In all 31 Delta sequences were submitted to GISAID. Mutations observed in different genes of Delta genome sequences are listed below with percentage of abundance of each mutation amongst the sequences. ORF1a includes A1306S (84%), T1395A (3%), P2046L(88%), P2287S (88%), A2529V(3%),V2930L(88%), T3255I(88%), L3606F(3%), T3646A (88%). ORF1b includes P314L (100%), G662S(100%), P1000L(100%), A1918V (88%). Spike region includes S13T (3%), T19R (28%), T95I (72%), Y145H (3%), E156- (100%), F157-(100%), R158G (100%), A222V (13%), L452R (94%), T478K (94%), D614G (94%), P681R (94%), D950N (97%), V1264L (3%). ORF3a includes S26L (97%). M gene includes I82T (97%), ORF7a includes V82A (94%) and T120I (97%) ORF7b includes T40I (88%), ORF8 includes Y42C (3%), D119-(38%) and F120-(100%). N gene includes D63G (100%), R203M (100%), G215C (88%) and D377Y (100%). ORF9b includes T60A (100%).

**Supplement Table S3:**

A total of 185 Omicron (5 Omicron BA.1+180 Omicron BA.2) sequences were submitted to GISAID. Mutations in different genes of Omicron genome sequences are listed below with percentage of abundance of each mutation amongst the submitted sequences. Mutations in ORF1a include S135R (98%), T842I (98%), G1307S (98%), R1421K (7%), L3027F (98%), T3090I (98%), L3201F (98%), T3255I (100%), P3395H (100%), S3675-(100%), G3676-(100%) and F3677-(98%). Mutations in ORF1b include P314L (100%), R1315C (98%), I1566V (100%), T2163I (98%) and S959P(44.3%).

Mutations in Spike include T19I (87%), L24-(97%), P25-(97%), P26-(97%), A27S (97%), G142D (97%), V213G (96%), G339D (100%), S371F (98%), S373P (100%), S375F (100%), T376A (98%), D405N (98%), R408S (98%), K417N (100%), S477N (100%), T478K (100%), E484A (100%), Q493R (100%), Q498R (100%), N501Y (100%),Y505H (100%), D614G (100%), H655Y (100%), N679K (100%), P681H (96%), N764K (100%), D796Y (100%), Q954H (100%) and N969K (100%). Mutations in ORF3a include T223I (97%). Mutations in E gene include T9I (98%) Mutations in M gene include Q19E (98%) and A63T (100%).Mutations in ORF6 include D61L (96%). Mutations in N gene include P13L (100%), E31-(100%), R32-(100%), S33-(100%), R203K (88%), G204R (100%) and S413R (97%). Mutations in ORF9b include P10S (100%), E27-(100%), N28 (100%)-and A29-(100%). Immune evasion mutation S371L present in Omicron BA.1 was substituted by S371F (98%), better immune evasion mutation in BA.2.^25^

## References

1. Ozono S, Zhang Y, Ode H, et al. SARS-CoV-2 D614G spike mutation increases entry efficiency with enhanced ACE2-binding affinity. Nat Commun. 2021;12(1):848. doi:10.1038/s41467-021-21118-2

2. Ramesh S, Govindarajulu M, Parise RS, et al. Emerging SARS-CoV-2 Variants: A Review of Its Mutations, Its Implications and Vaccine Efficacy. Vaccines. 2021;9(10). doi:10.3390/vaccines9101195

3. Jangra S, Ye C, Rathnasinghe R, et al. SARS-CoV-2 spike E484K mutation reduces antibody neutralisation. The Lancet Microbe. 2021;2(7):e283–e284. doi:10.1016/S2666-5247(21)00068-9

4. https://www.who.int/news/item/22-02-2022-statement-on-omicron-sublineage-ba.2. Statement on Omicron sublineage BA.2.

5. Yamasoba, Daichi, Izumi Kimura, Hesham Nasser, Yuhei Morioka, Naganori Nao, Jumpei Ito, Keiya Uriu et al. “Virological characteristics of SARS-CoV-2 BA. 2 variant.” BioRxiv (2022). doi:doi.org/10.1101/2022.02.14.480335

6. INSACOG WEEKLY BULLETIN, 10 January 2022 (https://dbtindia.gov.in/sites/default/files/INSACOG%20WEEKLY%20BULLETIN%2010-01-2022.pdf)

7. Ghoshal U, Garg A, Vasanth S, et al. Assessing a chip based rapid RTPCR test for SARS CoV-2 detection (TrueNat assay): A diagnostic accuracy study. Hasnain SE, ed. PLoS One. 2021;16(10):e0257834. doi:10.1371/journal.pone.0257834

8. Cepheid. 2020. Xpert Xpress SARS-CoV-2. (Package insert.) US Food and Drug Administration, Silver Spring M https://www.fda.gov/media/136314/download. A 8 A 2020. No Title.

9. Wu F, Zhao S, Yu B, et al. A new coronavirus associated with human respiratory disease in China. Nature. 2020;579(7798):265–269. doi:10.1038/s41586-020-2008-3

10. Rambaut A, Holmes EC, O’Toole Á, et al. A dynamic nomenclature proposal for SARS-CoV-2 lineages to assist genomic epidemiology. Nat Microbiol. 2020;5(11):1403–1407. doi:10.1038/s41564-020-0770-5

11. Trifinopoulos J, Nguyen L-T, von Haeseler A, Minh BQ. W-IQ-TREE: a fast online phylogenetic tool for maximum likelihood analysis. Nucleic Acids Res. 2016;44(W1):W232–5. doi:10.1093/nar/gkw256

12. Katoh K, Misawa K, Kuma K, Miyata T. MAFFT: a novel method for rapid multiple sequence alignment based on fast Fourier transform. Nucleic Acids Res. 2002;30(14):3059–3066. doi:10.1093/nar/gkf436

13. Hadfield J, Megill C, Bell SM, et al. Nextstrain: real-time tracking of pathogen evolution. Bioinformatics. 2018;34(23):4121–4123. doi:10.1093/bioinformatics/bty407

14. López JL, Lozano MJ, Fabre ML, Lagares A. Codon Usage Optimization in the Prokaryotic Tree of Life: How Synonymous Codons Are Differentially Selected in Sequence Domains with Different Expression Levels and Degrees of Conservation. MBio. 2020;11(4). doi:10.1128/mBio.00766-20

15. Davis JJ, Olsen GJ. Modal codon usage: assessing the typical codon usage of a genome. Mol Biol Evol. 2010;27(4):800–810. doi:10.1093/molbev/msp281

16. Sharp PM, Tuohy TM, Mosurski KR. Codon usage in yeast: cluster analysis clearly differentiates highly and lowly expressed genes. Nucleic Acids Res. 1986;14(13):5125–5143. doi:10.1093/nar/14.13.5125

17. 1999. PJ. Analysis of codon usage. PhD dissertation. University of Nottingham., Nottingham, England.

18. Uhlén M, Fagerberg L, Hallström BM, et al. Proteomics. Tissue-based map of the human proteome. Science. 2015;347(6220):1260419. doi:10.1126/science.1260419

19. Puigbò P, Bravo IG, Garcia-Vallve S. CAIcal: a combined set of tools to assess codon usage adaptation. Biol Direct. 2008;3:38. doi:10.1186/1745-6150-3-38

20. Silverj A, Rota-Stabelli O. On the correct interpretation of similarity index in codon usage studies: Comparison with four other metrics and implications for Zika and West Nile virus. Virus Res. 2020;286:198097. doi:10.1016/j.virusres.2020.198097

21. Zhou J, Zhang J, Sun D, et al. The distribution of synonymous codon choice in the translation initiation region of dengue virus. PLoS One. 2013;8(10):e77239. doi:10.1371/journal.pone.0077239

22. Liu L, Iketani S, Guo Y, et al. Striking antibody evasion manifested by the Omicron variant of SARS-CoV-2. Nature. 2022;602(7898):676–681. doi:10.1038/s41586-021-04388-0

23. Cui Z, Liu P, Wang N, et al. Structural and functional characterizations of infectivity and immune evasion of SARS-CoV-2 Omicron. Cell. 2022;185(5):860–871.e13. doi:10.1016/j.cell.2022.01.019

24. Chen J, Wei G-W. Omicron BA.2 (B.1.1.529.2): high potential to becoming the next dominating variant. ArXiv. Published online February 10, 2022. http://www.ncbi.nlm.nih.gov/pubmed/35169598

25. Iketani S, Liu L, Guo Y, et al. Antibody evasion properties of SARS-CoV-2 Omicron sublineages. Nature. 2022;604(7906):553–556. doi:10.1038/s41586-022-04594-4

26. Ogawa J, Zhu W, Tonnu N, et al. The D614G mutation in the SARS-CoV2 Spike protein increases infectivity in an ACE2 receptor dependent manner. bioRxiv Prepr Serv Biol. Published online July 22, 2020. doi:10.1101/2020.07.21.214932

27. Kumar S, Bansal K. Cross-sectional genomic perspective of epidemic waves of SARS-CoV-2: A pan India study. Virus Res. 2022;308:198642. doi:10.1016/j.virusres.2021.198642

28. Paz M, Aldunate F, Arce R, Ferreiro I, Cristina J. An evolutionary insight into Severe Acute Respiratory Syndrome Coronavirus 2 Omicron variant of concern. Virus Res. 2022;314:198753. doi:10.1016/j.virusres.2022.198753

29. van Dorp L, Richard D, Tan CCS, Shaw LP, Acman M, Balloux F. No evidence for increased transmissibility from recurrent mutations in SARS-CoV-2. Nat Commun. 2020;11(1):5986. doi:10.1038/s41467-020-19818-2

30. Simmonds P. Rampant C→U Hypermutation in the Genomes of SARS-CoV-2 and Other Coronaviruses: Causes and Consequences for Their Short-and Long-Term Evolutionary Trajectories. mSphere. 2020;5(3). doi:10.1128/mSphere.00408-20

31. Hui KPY, Ho JCW, Cheung M-C, et al. SARS-CoV-2 Omicron variant replication in human bronchus and lung ex vivo. Nature. 2022;603(7902):715–720. doi:10.1038/s41586-022-04479-6

32. Majumdar S, Sarkar R. Mutational and phylogenetic analyses of the two lineages of the Omicron variant. J Med Virol. 2022;94(5):1777–1779. doi:10.1002/jmv.27558

33. Kumar S, Thambiraja TS, Karuppanan K, Subramaniam G. Omicron and Delta variant of SARS-CoV-2: A comparative computational study of spike protein. J Med Virol. 2022;94(4):1641–1649. doi:10.1002/jmv.27526

34. Desingu PA, Nagarajan K. Omicron variant losing its critical mutations in the receptor-binding domain. J Med Virol. 2022;94(6):2365–2368. doi:10.1002/jmv.27667

35. Kannan, S.R.; Spratt, A.N.; Sharma, K.; Sönnerborg, A.; Apparsundaram, S.; Lorson, C.; Byrareddy, S.N.; Singh K. Complex Mutation Pattern of Omicron BA.2: Evading Antibodies without Losing Receptor Interactions. Preprints. 2022;(doi: 10.2. doi:10.20944/preprints202204.0120.v1

36. Chiara Pastorio, Fabian Zech Sabrina Noettger, Christoph Jung TJ, Kirchhoff KMJSF. Determinants of Spike Infectivity, Processing and Neutralization in SARS-CoV-2 Omicron subvariants BA.1 and BA.2. preprint. Published online 2022. doi:doi.org/10.1101/2022.04.13.488221

37. Lauring AS, Hodcroft EB. Genetic Variants of SARS-CoV-2-What Do They Mean? JAMA. 2021;325(6):529–531. doi:10.1001/jama.2020.27124

38. Weissman D, Alameh M-G, de Silva T, et al. D614G Spike Mutation Increases SARS CoV-2 Susceptibility to Neutralization. Cell Host Microbe. 2021;29(1):23–31.e4. doi:10.1016/j.chom.2020.11.012

