## Supplementary figures and images for "SARS-CoV-2 Omicron Variant Wave in India: Advent, Phylogeny and Evolution"

### Merged PDF Supplementary Figures.pdf

#### Delta, BA.1 and BA.2 lineages distribution

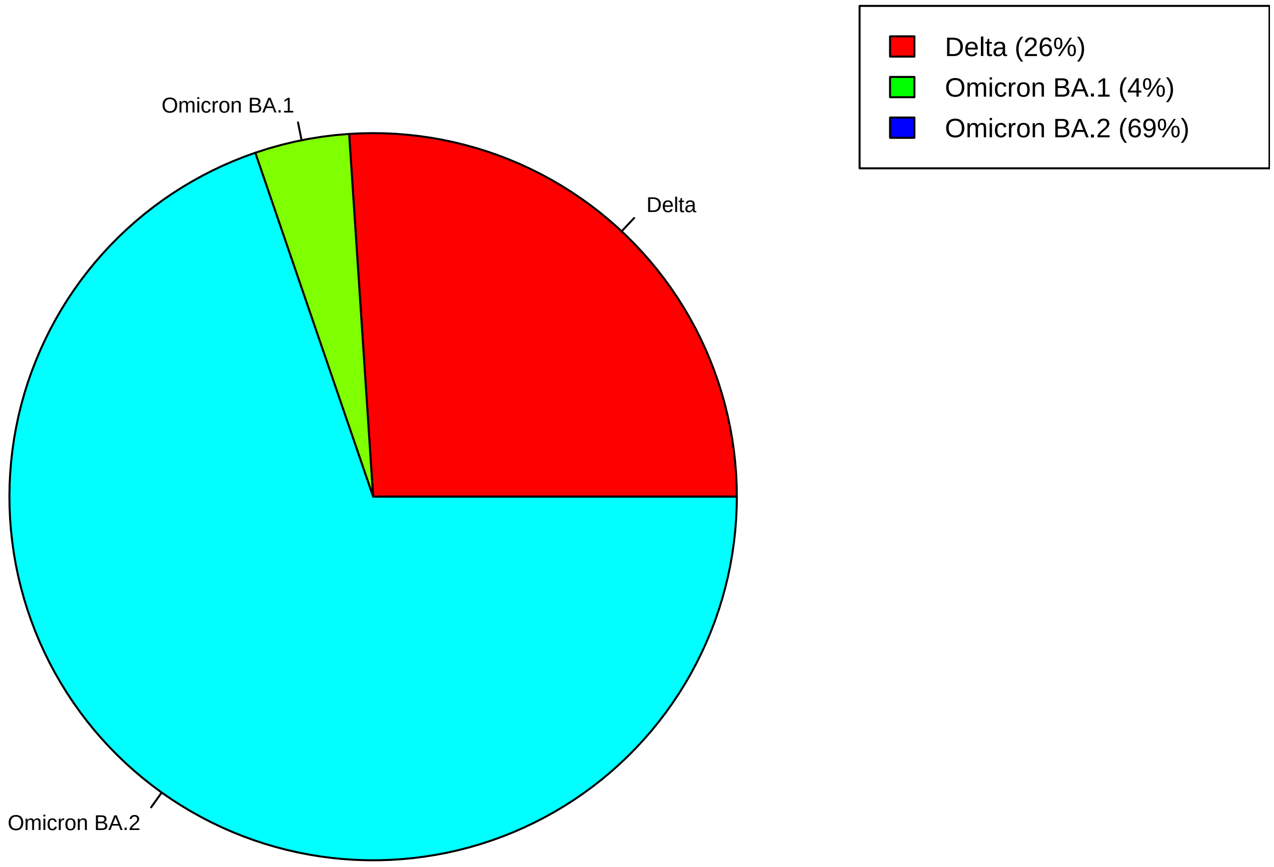


**Figure S1**

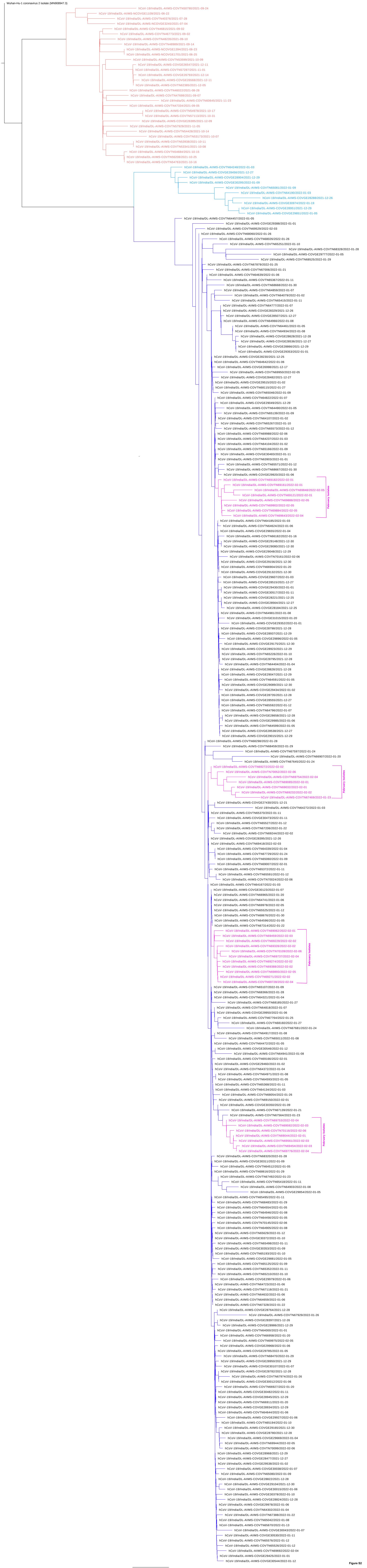

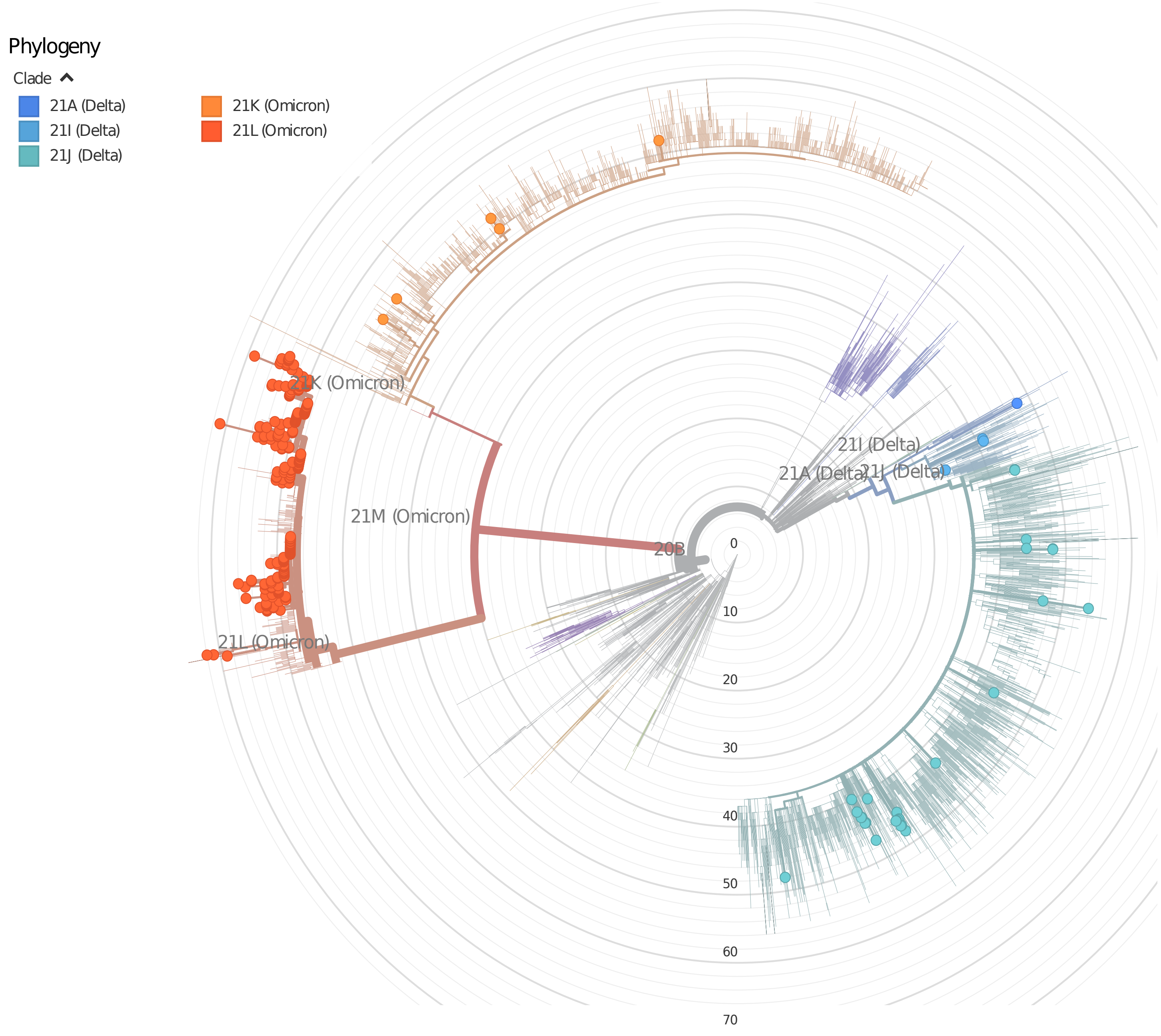

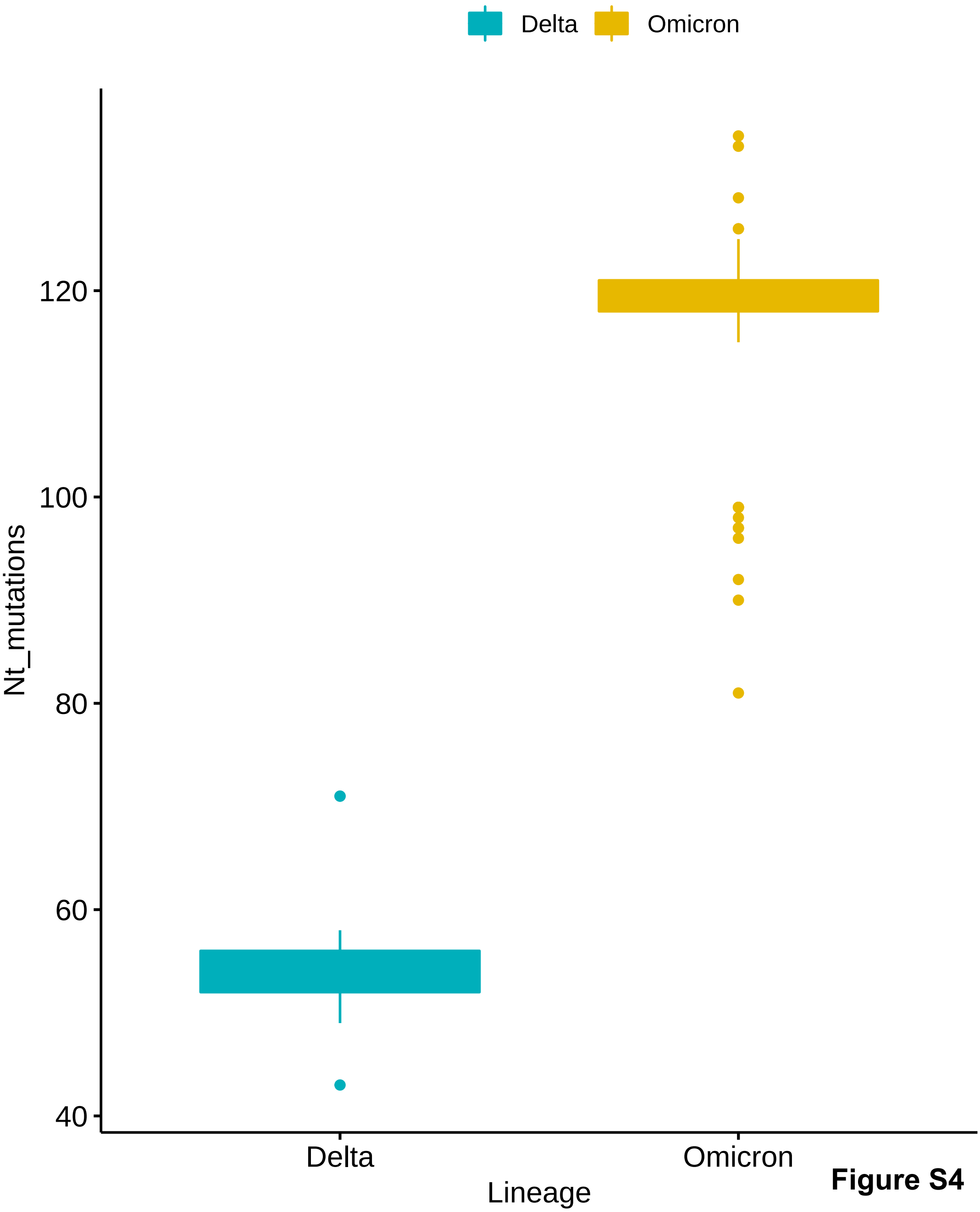
